# DUAL: deep unsupervised simultaneous simulation and denoising for cryo-electron tomography

**DOI:** 10.1101/2024.03.02.583135

**Authors:** Xiangrui Zeng, Yizhe Ding, Yueqian Zhang, Mostofa Rafid Uddin, Ali Dabouei, Min Xu

## Abstract

Recent biotechnological developments in cryo-electron tomography allow direct visualization of native sub-cellular structures with unprecedented details and provide essential information on protein functions/dysfunctions. Denoising can enhance the visualization of protein structures and distributions. Automatic annotation via data simulation can ameliorate the time-consuming manual labeling of large-scale datasets. Here, we combine the two major cryo-ET tasks together in DUAL, by a specific cyclic generative adversarial network with novel noise disentanglement. This enables end-to-end unsupervised learning that requires no labeled data for training. The denoising branch outperforms existing works and substantially improves downstream particle picking accuracy on benchmark datasets. The simulation branch provides learning-based cryo-ET simulation for the first time and generates synthetic tomograms indistinguishable from experimental ones. Through comprehensive evaluations, we showcase the effectiveness of DUAL in detecting macromolecular complexes across a wide range of molecular weights in experimental datasets. The versatility of DUAL is expected to empower cryo-ET researchers by improving visual interpretability, enhancing structural detection accuracy, expediting annotation processes, facilitating cross-domain model adaptability, and compensating for missing wedge artifacts. Our work represents a significant advancement in the unsupervised mining of protein structures in cryo-ET, offering a multifaceted tool that facilitates cryo-ET research.

## 1 Introduction

Cellular cryo-electron tomography (cryo-ET) is a powerful 3D imaging technique that visualizes sub-cellular structures at near-atomic resolution. Unlike single-particle cryo-EM, which isolates and purifies the target protein structure through biochemical means, cryo-ET directly images the complex sub-cellular structures in their native cytoplasm environment [1]. Given this unique advantage of preserving their spatial organization *in situ*, cryo-ET has been extensively applied to the study of the native structure, dynamic interaction, and spatial distribution of macromolecular complexes [2–4]. The 3D fine structures inside single cells provided by cryo-ET have potentially powerful applications in medical diagnostics as the dysfunctions of cellular structures may appear before any clinical symptoms [5–8]. Accordingly, interests in cryo-ET have been rapidly growing in the biomedical research community in recent years [9].

Sophisticated computational data processing methods are necessary to achieve the ultimate goal of visual proteomics: a complete structural description of the cell’s native molecular landscape [10]. Since 2017, as a result of the rapid development of deep learning techniques, supervised learning models have been proposed in cryo-ET for semantic segmentation [11,12], subtomogram classification [13], and object detection [14]. Nevertheless, a large amount of ground truth training labels, which mainly come from time-consuming annotation by a combination of traditional methods and manual selection, is required to attain the superior performance of supervised learning models. Researchers have tackled this new challenge by developing semi-supervised and unsupervised methods for tomogram segmentation [15], tomogram denoising [16,17], subtomogram alignment [18,19], and subtomogram clustering [20]. Despite being more sophisticated than supervised methods, unsupervised methods do not require labeled training data so that the labeling effort and subjective biases are considerably reduced. Consequently, with the fast development of unsupervised models in the computer vision field, they are expected to be applied for more and more tasks in cryo-ET.

To address the issues of data annotation and processing cost in two related major cryo-ET tasks, data simulation and denoising, we propose DUAL (Deep Unsupervised simultAneous denoising and simuLation) to combine them together in a single unsupervised framework. We have systematically evaluated DUAL on six datasets. Compared with popular denoising methods [16,21,22], DUAL achieved the best performance on the SHREC 2021 benchmark dataset [23] and in improving the particle picking accuracy on the *RELION* benchmark dataset [24]. For the tomogram simulation task, DUAL generated synthetic tomograms with indistinguishable styles, noise levels, and missing wedges to experimental tomograms. These realistic synthetic tomograms can be used to train semantic segmentation neural network models. When predicting on experimental tomograms, membranes and macromolecules of various molecular weights (from 560 to 3326 kDa), including ribosome, proteasome, TRiC, ClpB, and rubisco, are successfully detected and validated by subsequent subtomogram averaging. Furthermore, we have demonstrated other functionalities of DUAL: (1) DUAL can convert the styles between experimental tomograms of different imaging sources, such as low-SNR tomograms to high-SNR tomograms, for the purpose of noise reduction to desired levels, tomogram simulation with natural packing models, and domain adaptation for neural network training; and (2) DUAL can perform unsupervised learning based missing wedge compensation directly on the 3D reconstructed tomograms for reducing resolution anisotropy. The tutorial, code, and demo models will be available through the open-source Github software *AITom* [25] to provide easy and user-friendly access to the cryo-ET community.

## 2 Results

### 2.1 DUAL framework

In cryo-ET, individual 2D projection images are collected under an electron microscope by tilting the cellular specimen through a series of view angles. To prevent excessive electron beam damage to subsequent imaging at different angles, researchers usually set a low electron dosage and a limited tilt-angle range, resulting in the low Signal-to-Noise Ratio (SNR) and missing wedge effect of reconstructed 3D tomograms [26]. Therefore, advanced data processing techniques are required to assist researchers to interpret cryo-ET data both qualitatively and quantitatively. In this regard, traditional geometric or statistical methods have been proposed in cryo-ET including 3D reconstruction [27,28], missing wedge compensation [29,30], noise reduction [31,32], target macromolecule detection [33], membrane detection [34], subtomogram alignment [35,36], subtomogram classification and averaging [37–39], structural variability analysis [40,41], and tomogram data simulation [42,43].

Traditional geometric methods utilize pre-set rules and manually crafted features [44]. In contrast, supervised learning models optimize their massive parameters automatically through the guidance of training data labels. An important approach to reduce the dependency of deep learning models on labeled training data is through cryo-tomographic data simulation. Synthetic tomograms have pre-specified labels that can be used to test, benchmark, or tune relevant algorithms [45]. For example, the robustness of an analysis algorithm can be tested through performing on synthetic datasets with a range of imaging parameters such as spherical aberration, defocus, noise level, and missing wedge. Existing simulation methods generally consist of a projection and reconstruction model following a pre-processing packing model [46]. In the packing model, the structural density maps of target structures, including the cellular ultrastructure and embedding ice layers, are packed together into a 3D structural density map (a.k.a. grand model) to mimic the crowded cellular environment. In the projection and reconstruction model, the 3D structural density map is projected to 2D images and re-projected back with parameters simulating the actual tomographic imaging and reconstruction procedure. However, the existing algorithms [45–47] rely heavily on manually set parameters and certain assumptions, such as Gaussian white noise, to add the imaging artifacts and noises. These pre-defined factors and assumptions may produce unrealistic synthetic results. Utilizing the power of deep learning, it would be beneficial to automatically learn the characteristics of experimental data to simulate tomograms indistinguishable from experimental ones. With such realistically simulated tomograms, deep learning models can be trained directly and applied effectively to experimental data as a solution to the laborious training data annotation challenge.

A closely related cryo-ET task is denoising, which can be viewed as the inverse process of simulation. In simulation, a noise-free structural density map is translated into a noisy tomogram whereas in denoising, a noisy tomogram is translated into a noise-free structural density map. Due to the low SNR of tomograms, denoising facilitates the visualization and biological interpretation as well as downstream tasks such as particle picking [26], membrane detection [48], structural segmentation [49], and filament tracing [50]. Currently, there exist both traditional and deep learning based methods for cryo-ET denoising. Traditional methods employ carefully designed mathematical or statistical models, like wavelet-based filters [31] and Monte Carlo sampling [51], to enhance the structural signal. Meanwhile, deep learning based methods avoid modeling the noise pattern explicitly. Supervised approach requires carefully prepared ground truth denoised version of tomograms by averaging and aligning structures [52]. Unsupervised approaches have been proposed to learn from 2D projection images. *Topaz* [16] is trained with aligned and paired noisy 2D projection images. *SC-Net* [17] learns 3D denoising from filtered subsets of 2D projection images. Yet the complicated and time-consuming 3D reconstruction process may make this unsupervised approach less practical. So far, there is no unsupervised denoising approach to perform model training directly on 3D tomograms.

Inspired by the *CycleGAN* model [53], we propose DUAL (Figure 1), an unpaired image-to-image translation framework with a novel module to disentangle the noise latent factor from the underlying structure. From an image-to-image translation perspective, the simulation task is to translate a cryo-ET structural density map, generated from a packing model [23,54], into a synthetic tomogram whereas the denoising task is to translate a tomogram, collected experimentally, into a realistic structural density map. We denote the sample space of the structural density maps as the clean domain and the sample space of experimental tomograms as the noisy domain. Specifically, unlike most of image-to-image translation tasks [55], this task is asymmetric as there exists a one-to-many correspondence relationship between the clean domain and the noisy domain. A tomogram has only one corresponding structural density map as denoising ground truth. In contrast, given a structural density map, there is an infinite number of possible corresponding synthetic tomograms with different noises. Therefore, DUAL is designed to extract noise codes from the noisy domain and generate random noise codes to create random synthetic noises.

**Figure 1:**
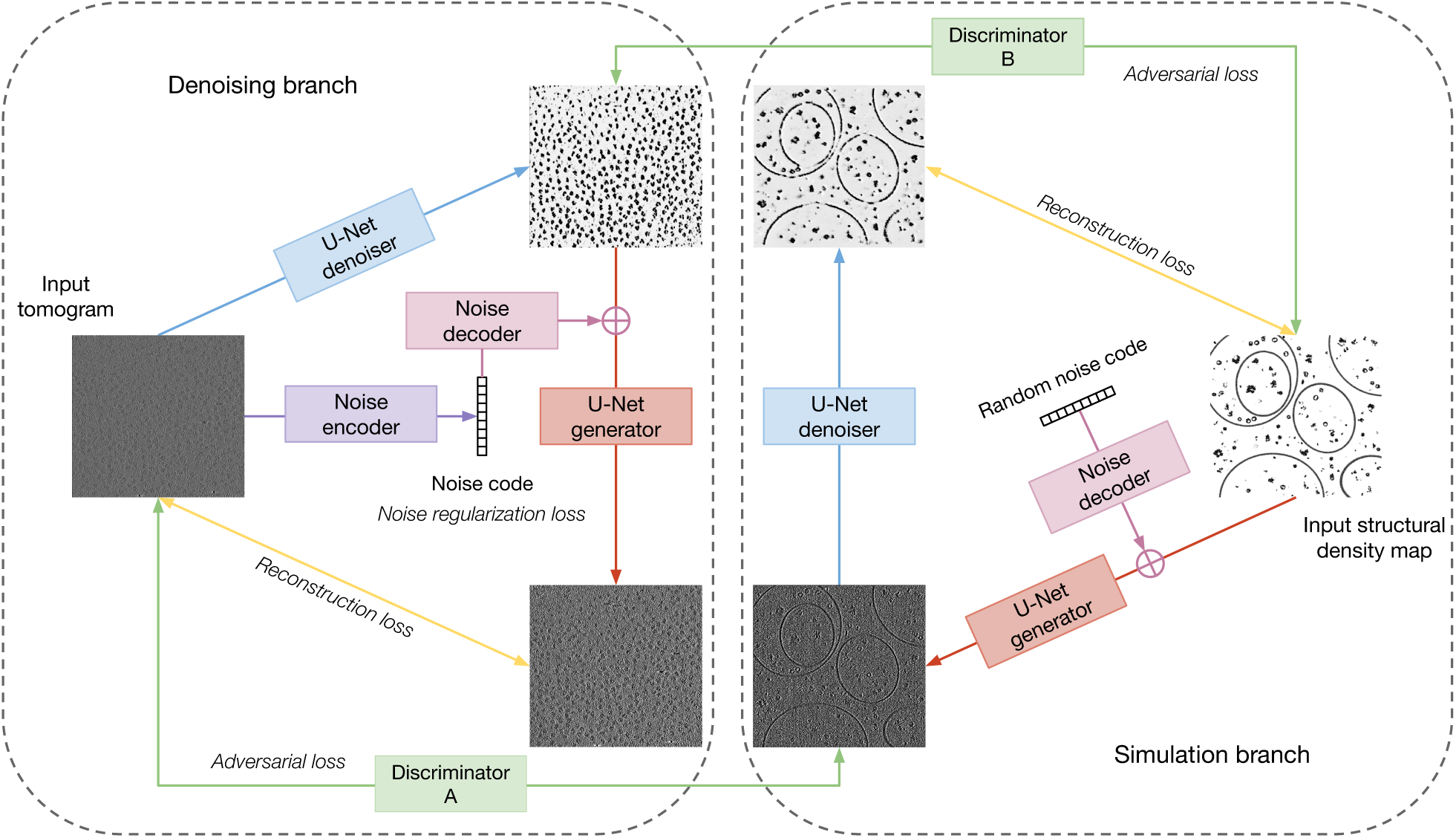
Conceptual workflow of DUAL. DUAL consists of six neural networks (detail architectures in supplementary note 1): U-Net denoiser, U-Net generator, noise encoder, noise decoder, and two discriminators. The inputs are a set of structural density maps from the clean domain and a set of tomograms from the noisy domain. We use two U-Nets [56] to translate images between the clean domain and the noisy domain. To address the one-to-many correspondence issue, we design a noise encoder to extract noise code from a noisy input and a noise decoder that could generate noise masks from noise codes. The noise decoder can take random noise codes to generate an infinite number of random noise masks for simulation. We employ discriminators [57] operating in both the spatial and the spectral space to learn the specific style of a domain in an adversarial fashion. In each epoch, the discriminators are trained to distinguish between real and fake images of a domain whereas the U-Net generators are trained with adversarial loss functions to “fool” the discriminators. In the simulation branch, the reconstruction loss function is used to preserve the structures. In the denoising branch, the reconstruction loss function and the noise regularization loss function are used together to correctly disentangle the noise pattern from structures. After training, the U-Net denoiser in the denoising branch can be deployed for tomogram denoising. Similarly, the U-Net generator and noise decoder in the simulation branch can be deployed for tomogram simulation.

### 2.2 Tomogram denoising

To evaluate the denoising performance of DUAL, we first applied it to the SHREC 2021 benchmark dataset. After training on the training dataset consisting of unpaired images from the noisy domain and the clean domain, the denoiser was applied to the testing tomogram (Figure 2A). We quantitatively evaluated DUAL and baselines (supplementary note 2) by measuring their Peak Signal-to-Noise Ratio (PSNR) and Structural Similarity Index Measure (SSIM) to the ground truth grand model. PSNR measures the ratio between the maximum power of structural signal and the power of noise that affects the fidelity of the denoised representation. The higher the PSNR, the higher the structural signal relative to noise. SSIM measures the preservation of structural information by focusing on strongly inter-dependant pixels, such as the edge of an object, to assess the denoising quality. SSIM ranges between 0 and 1. The higher the SSIM, the better the perceived structural information. Table 1 presents the PSNR and SSIM of denoised versions in reference to the ground truth. Compared to the tomogram without denoising (None), all methods showed some improvements in PSNR or SSIM, confirming that noises are partially reduced by these methods. DUAL achieves both the best SSIM and PSNR, confirming our qualitative observation that DUAL performs the best in reducing noise while preserving structural information. It can be observed in Figure 2A that NLM and Topaz-2 have relatively weaker noise reduction, which is reflected in their smaller change in PSNR. NAD and DUAL have relatively stronger noise reduction, which is also reflected in their larger improvements in PSNR. In cryo-ET, there is usually a trade-off between noise reduction and preservation of structural details, because both the noises and structural details exist mostly as the high-frequency components of the spectral domain. When reducing the noise, fine structural details may also be eliminated. SSIM is a more sophisticated metric which bases on three comparison functions of luminance, contrast, and structure in a small window size such as 7^3^. Without denoising, the SSIM is measured as 0.011, DUAL has a much larger improvement to 0.568 whereas all baselines have SSIM less than 0.1. Because the original tomogram, the ground truth, and all the denoised versions are standardized by subtracting their mean and dividing by their standard deviation when computing the SSIM, there should be little difference in their luminance comparison function. Therefore, the significant improvement in SSIM can be attributed to the better contrast between the structure and background as well as the finer structural shapes in each small window region.

**Figure 2:**
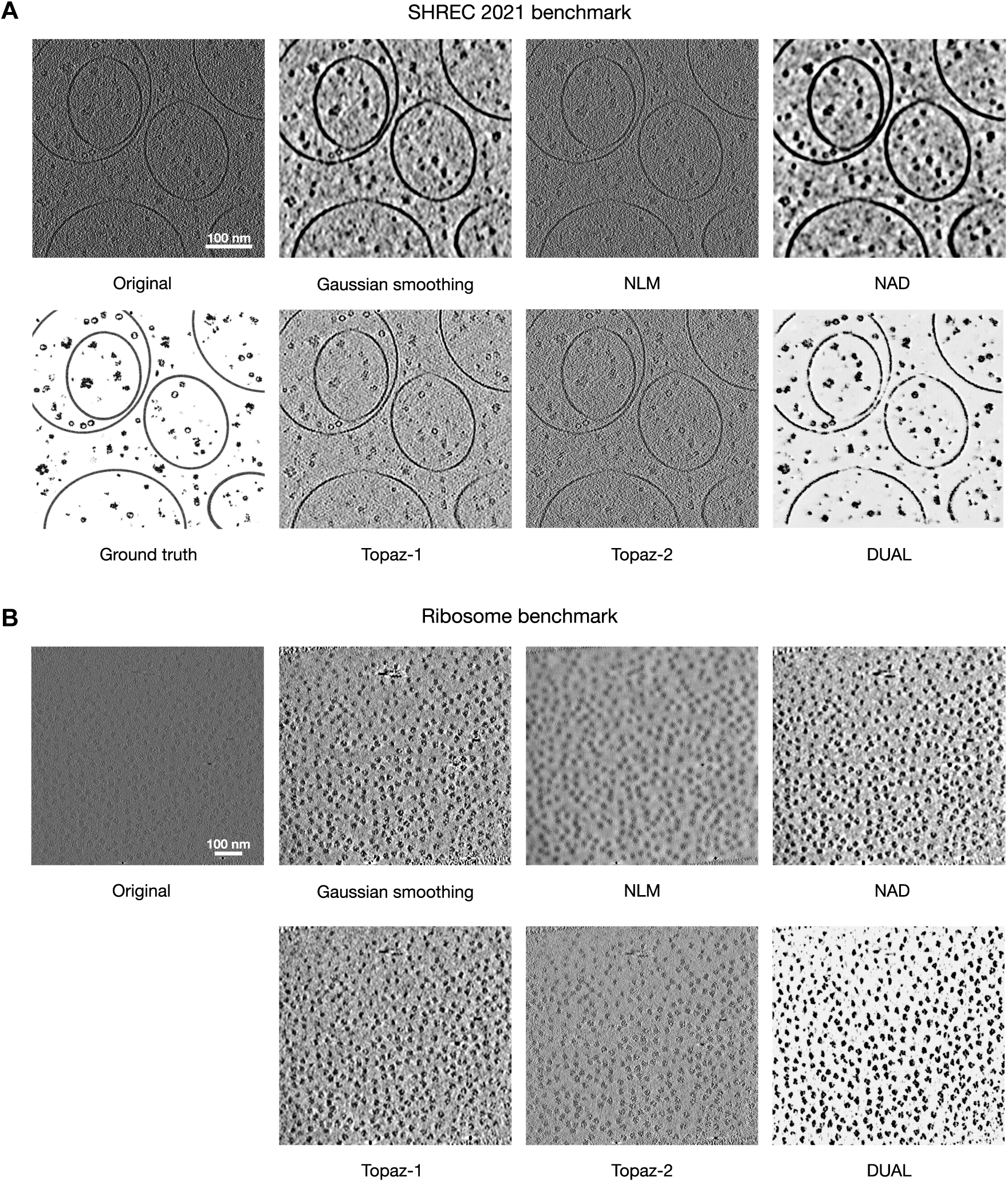
Tomogram denoising by DUAL and baseline methods. **A.** The original testing tomogram from the SHREC 2021 benchmark dataset, the grand model, and denoised versions by DUAL and four baseline methods. DUAL achieves visually cleaner results as indicated by the higher contrast between structure and background. The highest similarity between the DUAL denoised result and the ground truth evidences that DUAL provides the most effective noise reduction while preserving structural details. **B.** An example tomogram from the ribosome benchmark dataset and denoised versions by DUAL and four baseline methods. DUAL generates denoising results with the best contrast and visually clearest ribosome locations and shapes.

**Table 1:** Quantitative denoising evaluation on the SHREC 2021 benchmark dataset.

| Methods | PSNR | SSIM |
| --- | --- | --- |
| None | 32.33 | 0.011 |
| Gaussian smoothing | 34.93 | 0.085 |
| NAD | 36.06 | 0.097 |
| NLM | 32.24 | 0.012 |
| Topaz-1 | 34.48 | 0.057 |
| Topaz-2 | 32.65 | 0.019 |
| DUAL | **37.01** | **0.568** |

We then evaluated the denoising performance of DUAL on the experimental ribosome benchmark dataset (Figure 2B).

Unlike synthetic datasets, experimental datasets do not have the ground truth of structural density maps for quantitative comparison. Therefore, we evaluated the denoising performance on experimental datasets using indicators from downstream tasks. One important goal of cryo-ET denoising is to improve the downstream particle picking accuracy. As better denoising generally leads to more accurate particle picking, we utilized particle picking accuracy as the indicator of denoising performance. Tomograms in the ribosome benchmark dataset contain isolated and purified 80S ribosome complexes. The particle location ground truth has been provided by the authors through manual picking [24]. We applied a popular template-free particle picking method, Difference of Gaussians (DoG), to the tomograms and their denoised versions by each method. We controlled the hyper-parameters to be the same in all experiments to pick the top 500 detections in each tomogram. Due to the fact that the diameter of a yeast ribosome is around 28 nm, any DoG detection within 8 nm distance of a ground truth particle location is considered an overlap and counted as a true positive. The results are summarized in Table 2. The precision measures the percentage of DoG detections that overlaps with the ground truth locations. The recall measures the percentage of ground truth locations that have DoG detections overlapping with them. The F1 score is the harmonic mean of precision and recall to provide a balance between them. DUAL had significant improvement in particle picking precision, recall, and F1 score over baseline methods. Applying DoG directly to the tomograms without denoising resulted in an average F1 score of 0.577. Applying DoG on tomograms denoised by baseline methods results in average F1 scores ranging from 0.495 to 0.586, which shows only marginal improvements. The F1 score after DUAL denoising is significantly improved to 0.702. The standard deviations in the precision, recall, and F1 score of DUAL are also lower than those of baseline methods, suggesting that the denoising performance of DUAL is stable and consistent in improving the particle picking results across different tomograms. The deep learning based *Topaz* model [16] is a Noise2Noise framework that trains on paired observations to minimize the L2 reconstruction error across them. Models based on the Noise2Noise framework [58] require carefully prepared paired observations for training, whereas DUAL only requires samples from an unpaired clean domain such as publicly available structural density maps. The outperformance of DUAL to Topaz is likely due to the adversarial training of DUAL that effectively recognizes macro-molecular structures and hence successfully enhances the signal of ribosomes in this dataset.

**Table 2:** Particle picking accuracy on denoised ribosome benchmark dataset. Each cell contains the mean and standard deviation of the corresponding statistic across seven tomograms.

| Methods | Precision | Recall | F1 |
| --- | --- | --- | --- |
| None | $0.525 \pm 0.098$ | $0.645 \pm 0.134$ | $0.577 \pm 0.110$ |
| Gaussian smoothing | $0.535 \pm 0.092$ | $0.654 \pm 0.125$ | $0.586 \pm 0.101$ |
| NAD | $0.530 \pm 0.092$ | $0.648 \pm 0.126$ | $0.581 \pm 0.102$ |
| NLM | $0.453 \pm 0.085$ | $0.550 \pm 0.120$ | $0.495 \pm 0.097$ |
| Topaz-1 | $0.476 \pm 0.125$ | $0.581 \pm 0.122$ | $0.521 \pm 0.143$ |
| Topaz-2 | $0.513 \pm 0.103$ | $0.632 \pm 0.147$ | $0.564 \pm 0.119$ |
| DUAL | **$0.641 \pm 0.064$** | **$0.783 \pm 0.044$** | **$0.702 \pm 0.038$** |

### 2.3 Tomogram simulation

As a multi-task model, the simulation branch of DUAL is equally important as the denoising branch. Experimental tomograms are usually characterized by their low SNRs and missing wedge effects. To investigate whether the noise level and missing wedge effects are properly learned, we visualize in Figure 3B the synthetic tomograms simulated using the DUAL U-Net generators and noise decoders trained on the four datasets. Existing cryo-ET simulation works [23,45] assume Gaussian white noise and require the SNR and tilt-angle range to be set manually. DUAL is the first cryo-ET simulation framework that can automatically learn the noise pattern, noise level, and missing wedge effect through adversarial training to provide the most realistic synthetic results.

**Figure 3:**
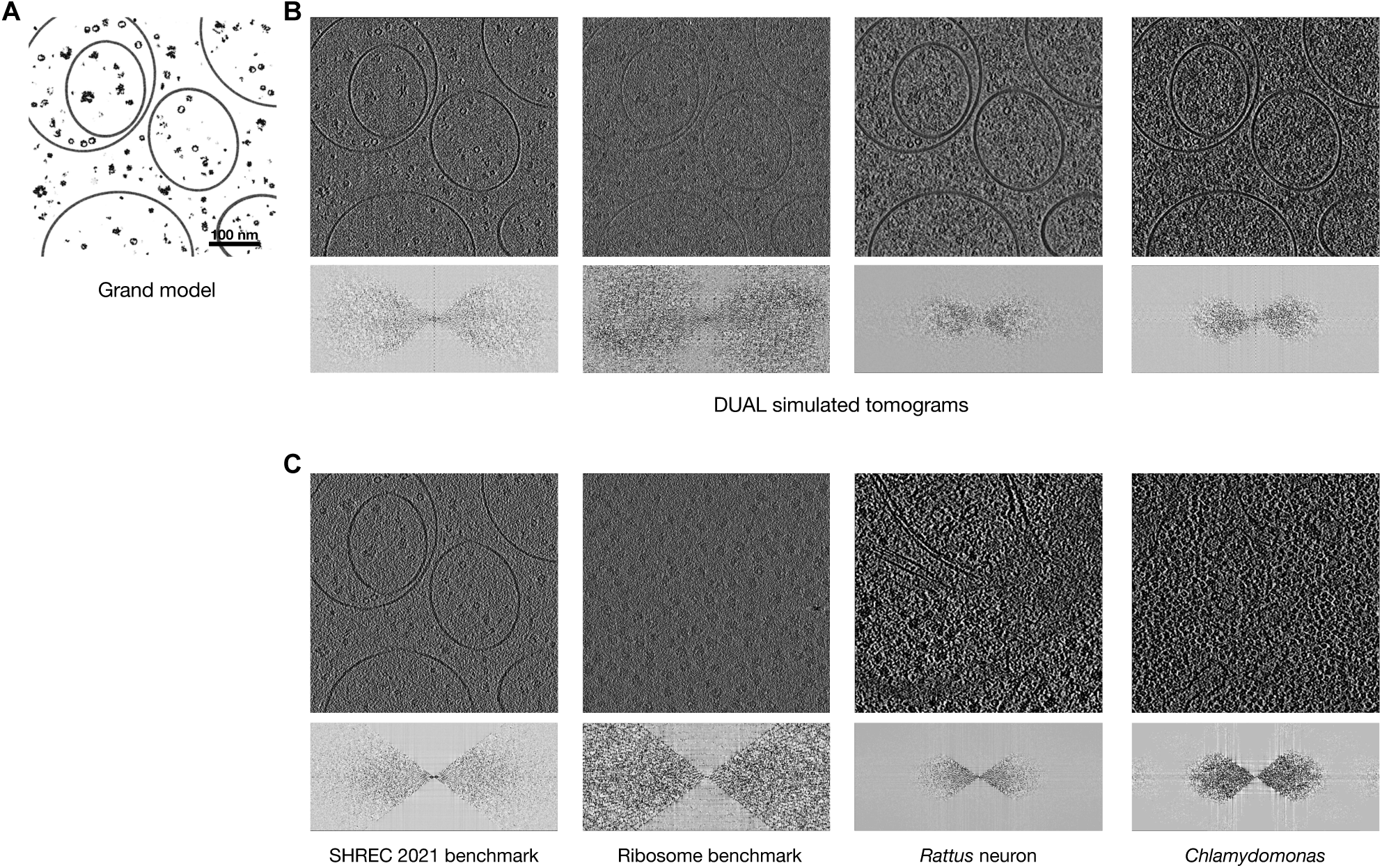
Tomogram simulation by DUAL. **A.** The grand model from the SHREC 2021 benchmark dataset. **B.** Synthetic tomograms simulated by DUAL learning from the cryo-tomographic styles of SHREC 2021 benchmark dataset, ribosome benchmark dataset, *Rattus* neuron dataset, and *Chlamydomonas* pyrenoid dataset. The Fourier space representations showing missing wedge effects are visualized below each tomogram. **C.** Original tomograms from corresponding dataset. The synthetic tomograms have visually similar noise levels and noise patterns to their corresponding experimental tomograms. The similar noise level and pattern can also be validated by the visualization of the spectral representation. The high-frequency components are usually dominated by noises. For example, the ribosome benchmark has a higher level of noise and therefore more high-frequency signals. The synthetic tomogram trained using the ribosome benchmark dataset also has more high-frequency components. In addition, the spectral representations show that DUAL has successfully learned the missing wedge patterns of different datasets.

Similar to the denoising evaluation, we measure the downstream task performance as an indicator of the simulation performance. A major goal of realistically simulated tomograms is to provide training data with readily available pre-specified labels for neural network training. The trained models can then be applied to predict on experimental tomograms as a transfer learning approach to reduce the training data annotation burden. Generally, the more similar the synthetic data to the experimental data, the better the prediction results are.

Since there are 16 semantic classes in the SHREC 2021 benchmark dataset, we select classes with significant abundance for visualization in Figure 4 and further subtomogram averaging analysis (supplementary Figure S3-S7). Through manual selection and subtomogram classification, the authors of the *Rattus* neuron dataset have discovered and recovered three macromolecular complexes: ribosome, TRiC/CCT chaperonin, and 26S proteasome. Based on their observations, the authors have concluded that neuronal poly-Gly-Ala aggregates recruit 26S proteasomes and exclude other large macromolecular complexes such as ribosomes and TRiC/CCT chaperonins [59]. Our DUAL simulation-based transfer learning semantic segmentation approach successfully segmented out the membrane structure and detected four macromolecular complexes. We not only validated the original authors’ detection of ribosome, TRiC/CCT chaperonin, and 26S proteasome, but also detected a new ClpB-like structure. ClpB (Caseinolytic peptidase B protein homolog) is a AAA ATPase chaperone that exists in the mitochondria. As shown in Figure 4B, the majority of ClpB-like structures (pink) are detected inside the mitochondria. Furthermore, the detected macromolecular structures are confirmed by subtomogram averaging with resolution < 32Å for effective recognition. The authors of the *Chlamydomonas* pyrenoid dataset [11] have developed a supervised semantic segmentation approach with manually prepared data annotation for training to detect rubisco holoenzymes. They have also manually segmented the pyrenoid tubule membranes. Using our DUAL simulation-based transfer learning approach, the pyrenoid tubule membranes and rubisco holoenzymes can be automatically segmented out. In comparison, if we train the semantic segmentation neural network model using the synthetic tomograms provided in the SHREC 2021 benchmark dataset, the membrane structure can be segmented out relatively successfully but most of the rubisco holoenzymes were misclassified to other macromolecular classes (Figure 4F). This demonstrates that DUAL generated better synthetic tomograms than traditional cryo-ET simulation approaches with manually set imaging parameters and additive Gaussian white noise. The realistic synthetic tomograms can be used to effectively facilitate downstream tasks such as the training of neural network models.

**Figure 4:**
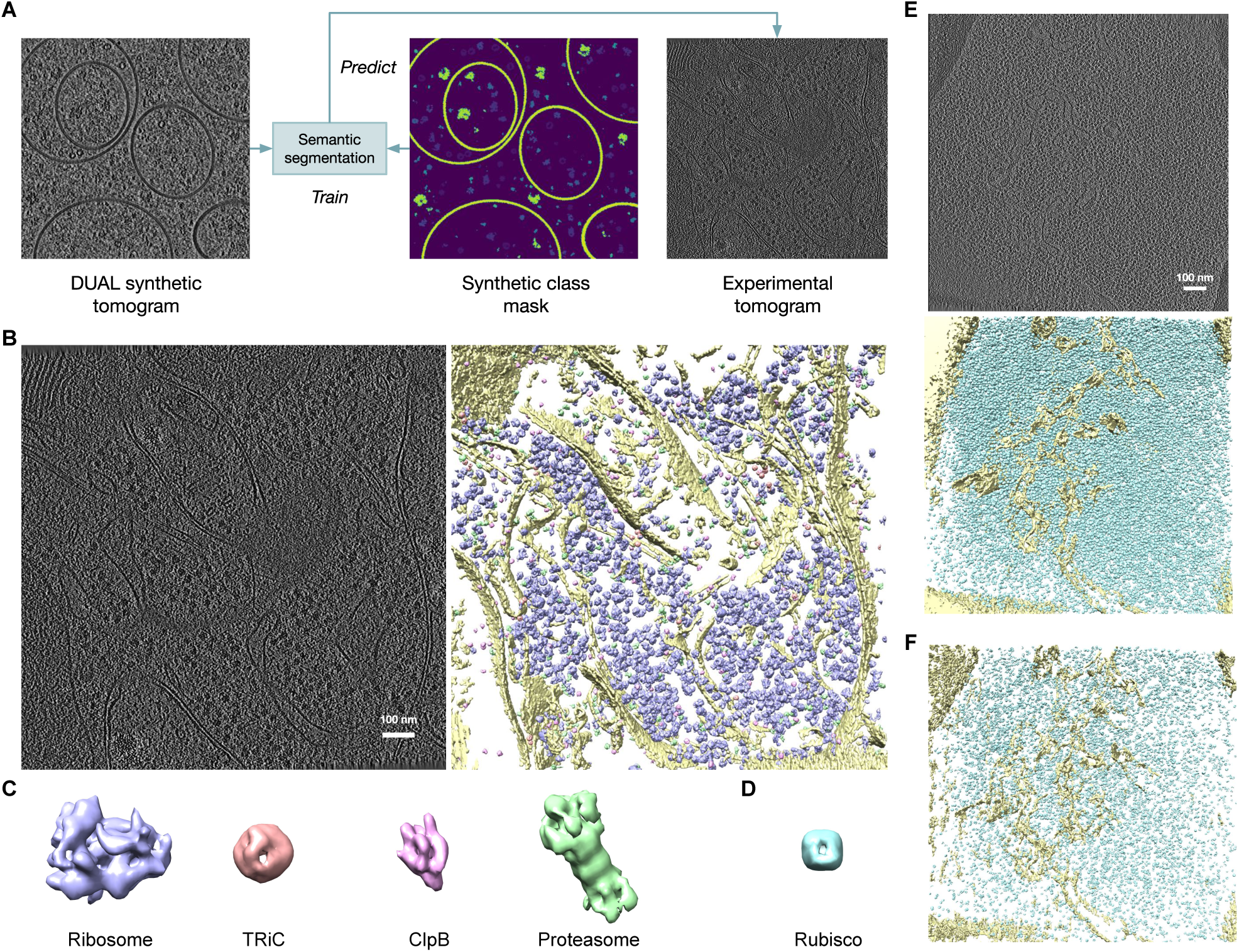
DUAL simulation-based transfer learning approach for semantic segmentation on experimental tomograms. **A.** Workflow: we first generated synthetic tomograms by applying the U-Net generators and noise decoders in DUAL to the grand models in the SHREC 2021 benchmark dataset. The DUAL models were trained using the *Rattus* neuron dataset and the *Chlamydomonas* pyrenoid dataset. Next, semantic segmentation neural network models, employing the network proposed in *Deepfinder* [11], were trained using the DUAL synthetic tomograms and segmentation ground truth masks in the SHREC 2021 benchmark dataset. Then the trained semantic segmentation neural network models were applied to the *Rattus* neuron dataset and the *Chlamydomonas* pyrenoid dataset, respectively. **B.** An example tomogram from the *Rattus* neuron dataset and corresponding iso-surface representation of 3D semantic segmentation of membrane structure (yellow), ribosome (indigo), TRiC (red), ClpB (pink), and 26S proteasome (green). **C.** Subtomogram averages of detected macromolecular complexes. **D.** Subtomogram average of detected rubisco structure. **E.** The tomogram from the *Chlamydomonas* pyrenoid dataset and corresponding iso-surface representation of 3D semantic segmentation of membrane structure (yellow) and rubisco (blue). **F.** 3D semantic segmentation on the *Chlamydomonas* pyrenoid dataset with neural network model trained on the SHREC 2021 benchmark tomograms.

### 2.4 Clean domain construction

In the experiments above, we used the grand models (noise-free 3D structural density maps) provided in the SHERC 2021 benchmark dataset to construct the clean domain. We note that it is also possible to construct a clean domain from experimental tomograms with relatively high SNR. In this way, the low-SNR experimental tomograms in the noisy domain can be converted to high-SNR representations indistinguishable from the clean domain experimental tomograms, and *vice versa*. We conduct experiments using the low-SNR tomograms from the ribosome benchmark dataset as the noisy domain and the high-SNR tomograms from the *Chlamydomonas* chloroplast dataset and the SARS-CoV-2 infection dataset as two clean domains.

Constructing the clean domain with high-SNR experimental tomograms comes with two potential advantages. First, in the conventional simulation approach, grand models are generated by packing 3D structural densities together with manually set distributions. For example, the 13 types of macromolecular complexes in the SHREC 2021 benchmark dataset [23] are manually chosen and assumed to exhibit similar abundance and be distributed randomly according to 3D uniform distributions. Such packing models differ from the actual structural distributions and interactions in experimental data. Instead, if we simulate synthetic tomograms by generating low-SNR representations of high-SNR experimental tomograms, the natural structural packing in high-SNR tomograms will provide biologically plausible spatial organizations of structures. Second, this enables another potential learning-based semantic segmentation approach. If a set of experimental tomograms have available segmentation masks (preferably high-SNR ones as they are easily hand-segmented or ones obtained with fluorescence labeling through cryo-CLEM [60]), neural network models can be trained on this dataset. Then, another experimental dataset can be adapted to the high/low-SNR domain using DUAL and segmented using the trained semantic segmentation neural network model. As shown in Figure 5, we converted the experimental tomogram from the *Rattus* neuron dataset to its low-SNR representation using the tomograms in the SHREC 2021 benchmark dataset to construct the noisy domain. Then, the semantic segmentation neural network trained on the SHREC 2021 benchmark dataset was applied to the low-SNR representation of that experimental tomogram. We obtained similar semantic segmentation results (Figure 5C) to the one shown in Figure 4B. We note that the semantic segmentation of macromolecular complexes in Figure 5C is visually less clear compared to that of Figure 4B. This is likely due to the structural information loss during the conversion as the neural network is applied directly to the experimental tomogram in the DUAL simulation-based approach whereas the neural network is applied to the converted experimental tomogram in this domain adaptation approach. Therefore, the domain adaptation approach may be a sub-optimal transfer learning solution to cryo-ET semantic segmentation compared to the simulation-based approach. Nevertheless, the DUAL domain adaptation approach has the advantage of being more efficient. Only one semantic segmentation neural network needs to be kept rather than training a separate network for each synthetic dataset in the simulation-based approach. In brief, DUAL is essentially a flexible framework that can adapt diverse modalities for different biological meaningful functionalities.

**Figure 5:**
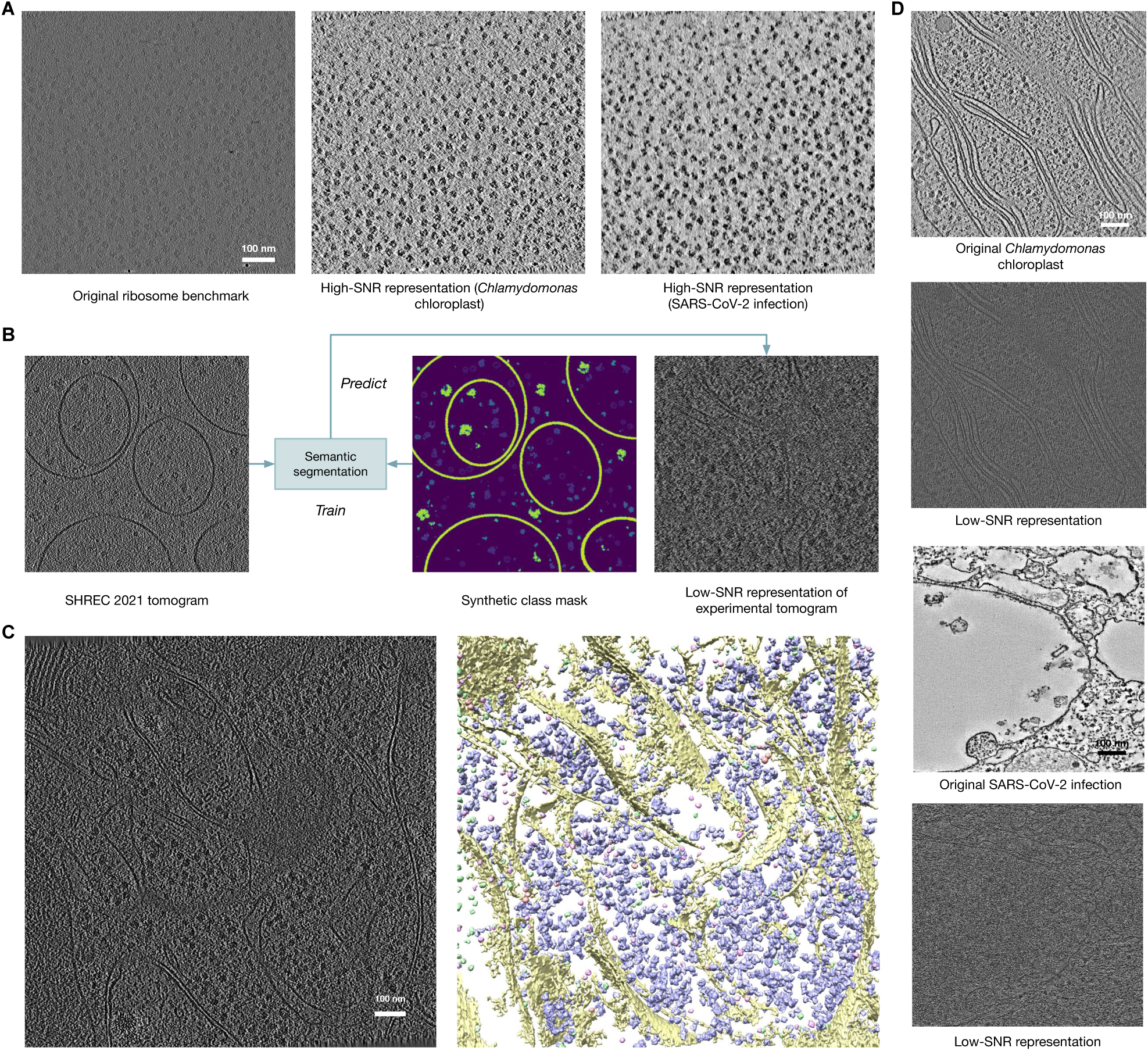
Clean domain constructed by high-SNR experimental tomograms. **A.** An example tomogram from the ribosome benchmark dataset and its high-SNR representations converted using DUAL U-Net denoiser. **B.** DUAL domain adaptation transfer learning approach for semantic segmentation on experimental tomograms. **C.** The example tomogram from the *Rattus* neuron dataset and corresponding iso-surface representation of 3D semantic segmentation. **D.** Example tomograms from the *Chlamydomonas* chloroplast dataset and SARS-CoV-2 infection dataset and their corresponding low-SNR representations converted using DUAL U-Net generator and noise decoder. DUAL can effectively convert the noisy tomogram from the ribosome benchmark dataset to its high-SNR representations and convert the relatively clean tomograms to their low-SNR representations. Both the high-SNR and low-SNR representations are visually similar to the style of their corresponding experimental tomograms. Therefore, desired noise reduction levels can be achieved through the choice of high-SNR experimental datasets for the clean domain.

### 2.5 Missing wedge compensation

Due to increases in effective thickness of the imaging sample at higher tilt angles, the tilt-angle range is typically limited to ±60° to prevent excessive radiation damage. This will result in the missing wedge effect which causes severe artifacts in the reconstructed tomogram such as distortion and elongation of sub-cellular structures [30]. The missing wedge effect hinders visual interpretation and subtomogram averaging, which is key to the analysis of macromolecular structures and spatial organizations *in situ*. Missing wedge compensation is a very challenging task in cryo-ET as the missing information in the spectral domain needs to be imputed. Existing works [29,30,61] proposed to compensate the missing wedge through *priori* assumptions during 3D reconstruction. A recent work, IsoNet [62], has pointed out the limitation of these existing works and proposed an unsupervised learning-based method to perform missing wedge compensation directly on 3D reconstructed tomograms.

Here, we show that the DUAL framework can be extended according to Figure 6A to perform missing wedge compensation directly on 3D reconstructed tomograms. We evaluated the performance of missing wedge compensation on the low-SNR ribosome benchmark dataset. As shown in Figure 6B, the missing cone region is clearly visible in uncompensated tomograms. Comparatively, the high-frequency region of the missing cone regions are filled more by DUAL than IsoNet. Since there is no ground truth for the missing information, to quantitatively evaluate the performance of the two methods, we first compensated the missing wedge (±60° along the z light-axis) on a tomogram by both methods. Then, we artificially created a non-overlapping missing wedge along the x light-axis by masking out the information not in the ±60° tilt-angle range from the Fourier space. On the tomogram with the artificial missing wedge, we performed missing wedge compensation again by each method. The compensated information can be quantitatively compared with the ground truth that was masked out.

**Figure 6:**
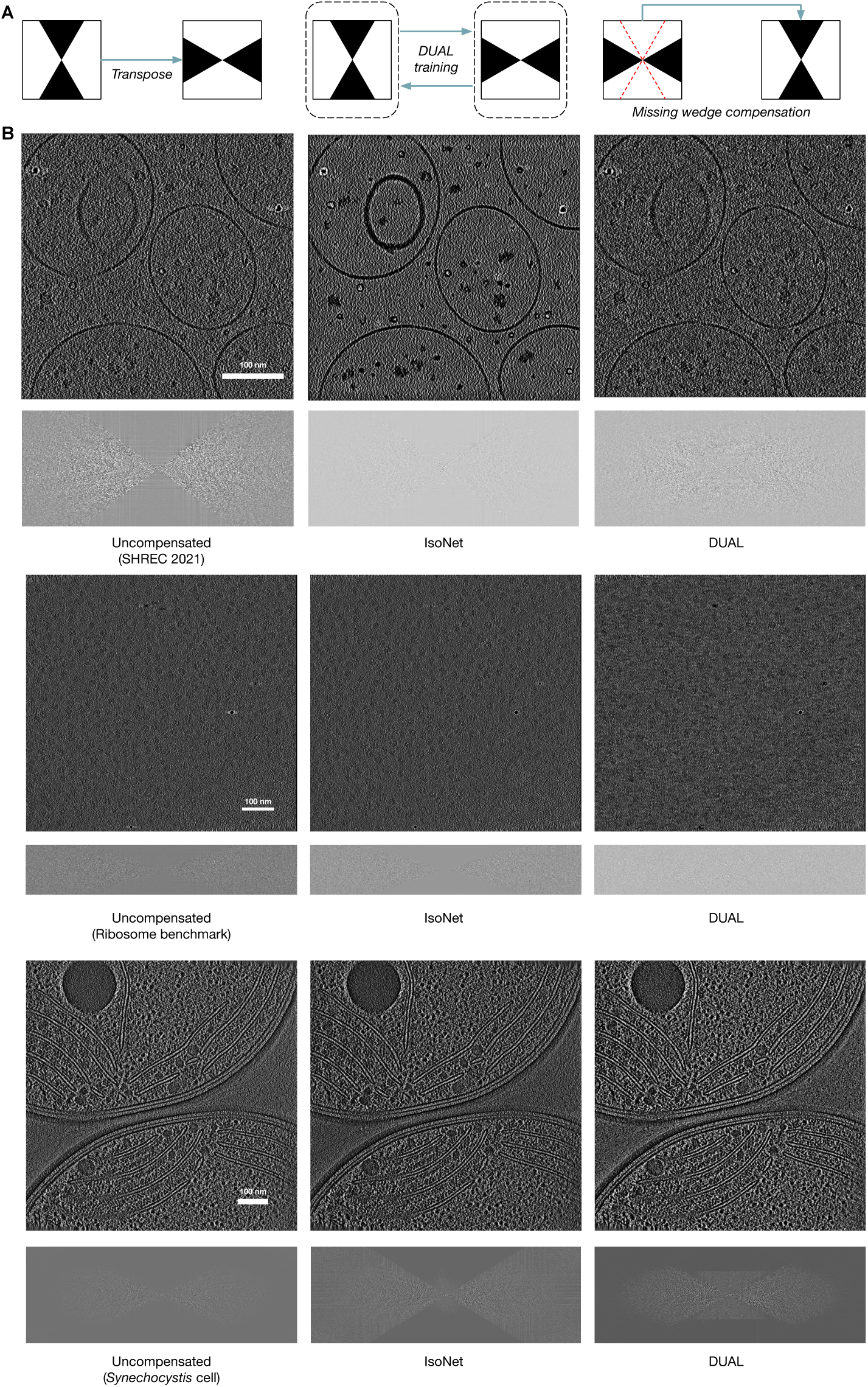
Missing wedge compensation using DUAL. **A.** We represent each tomogram as a mask of observed signals in the Fourier space and illustrate the workflow. Given a set of tomograms, for example, with tilt-angle range ±60°, the y-axis as the tilt-axis and the z-axis as the light axis, we could construct one domain using the original tomograms. Then, we could transpose the tomograms to construct another domain such that the y-axis remains the tilt axis and the x-axis becomes the light axis. The missing cone regions in the Fourier space of the two domains are non-overlapping. After unpaired image-to-image translation between these two domains using DUAL, the missing wedge effect in the original tomograms can be compensated by the information from the translated domain. **B.** Performance of DUAL and IsoNet [62] on three datasets.

As shown in Table 3, both methods achieved similar performance. Since DUAL has more high-frequency information compensated, it can be potentially used to complement IsoNet in missing wedge compensation. Consequently, missing wedge compensation using DUAL can facilitate the systematic analysis of cryo-ET data with improved imaging limits for biological discoveries *in situ*.

**Table 3:** Missing wedge compensation performance on three demo datasets. Each cell contains the PSNR and SSIM as reconstruction similarity measure. We note that since each method was performed twice for each tomogram in our evaluation, the results are not indicative for a one-time performance.

|  | SHREC 2021 | Ribosome benchmark | *Synechocystis* cell |
| --- | --- | --- | --- |
| IsoNet | 33.27, 0.108 | 37.99, 0.118 | 29.40, 0.054 |
| DUAL | 33.63, 0.147 | 35.97, 0.146 | 28.89, 0.043 |

## 3 Discussion

Cryo-electron tomography (cryo-ET) stands as a crucial method for precisely visualizing native sub-cellular structures at high resolution, offering immense potential that can be harnessed through advanced data processing techniques. However, several bottlenecks in cryo-ET data analysis and interpretation persist. The low signal-to-noise ratio (SNR) limits the ability to identify protein structures accurately and infer their functions or dysfunctions. Algorithms less robust to noise often struggle with the highly noisy nature of cryo-ET data. Additionally, the vast amount of three-dimensional data imposes high time costs on researchers for manual assessments and annotations, hindering quantitative evaluations. Deep learning approaches, while providing high-throughput automatic annotation, still require annotated data for effective model training.

This paper introduces DUAL, an innovative end-to-end unsupervised deep learning framework that simultaneously addresses two critical challenges in cryo-ET: denoising and data simulation. Leveraging a cyclic generative adversarial network with noise disentanglement, DUAL establishes an effective framework with unpaired training criteria. This framework translates between a noisy domain, comprising low-SNR tomograms, and a clean domain consisting of noise-free structural density maps (or high-SNR tomograms). Notably, DUAL achieves unsupervised cryo-ET denoising without relying on the sophisticated use of 2D projection images, marking a significant advancement. Simultaneously, it pioneers learning-based cryo-ET data simulation, generating synthetic tomograms with styles indistinguishable from experimental ones.

Our evaluation on the SHREC 2021 benchmark dataset showcases that the denoising branch of DUAL outperforms popular cryo-ET denoising methods, as evidenced by both peak signal-to-noise ratio and structural similarity index metrics. This noise reduction capability significantly enhances downstream tasks such as particle picking. The simulation branch of DUAL autonomously learns noise characteristics and missing wedge effects of experimental tomograms, producing highly realistic synthetic cryo-ET data. Importantly, this data can be efficiently employed without manual annotation to train neural network models for tasks like semantic segmentation, providing biologically valid results.

However, DUAL has its limitations. The reliance of the simulation branch on a pre-processing packing model introduces potential unnatural structural distributions, overlooking dynamic protein interactions. This limitation can be mitigated through the adoption of data-driven packing models or constructing the clean domain from high-SNR experimental tomograms with natural packing. Another limitation pertains to the level of noise reduction, as controlling noise reduction to preserve fine structural details remains a challenge. While the use of higher-SNR experimental data in DUAL partially addresses this, future work could focus on developing unsupervised learning-based cryo-ET denoising models with direct control over the level of denoising.

In summary, DUAL is a practical fully unsupervised multi-task learning framework, empowering cryo-ET researchers in various aspects. It enhances the visualization and annotation of sub-cellular structures, aids in accurate structure segmentation and template matching, benchmarks algorithms using simulated data, facilitates neural network model training using realistically generated synthetic data, verifies biological findings on low-SNR data by converting to high-SNR representations, and simplifies the cryo-ET imaging process. DUAL, characterized by its efficiency, completes training in only a few hours with GPU support, providing a potent alternative to complement existing cryo-ET data analysis approaches. By offering a powerful suite of functionalities, DUAL opens new opportunities for important discoveries in the structural biology community.

## Methods

### Model formulation

DUAL achieves unpaired/unsupervised training by adversarial learning. In paired image-to-image translation training, for each image, the ground truth translation target image is required for learning their correspondence relationship [52]. In comparison, unpaired image inputs from the two domains are sufficient in the unpaired/unsupervised setting. While the availability of paired tomogram and structural density map datasets is limiting supervised learning models from being widely adopted, unpaired training can facilitate denoising and simulating tomograms by focusing on the characteristics of the two domains rather than two paired images in order to create more generalizable models. Besides, the introduction of the noise disentanglement module enables DUAL to disentangle out and learn the noise pattern separately from structures, so as to provide more realistic synthetic data for downstream tasks.

Given a set of experimental tomograms sampled from the noisy domain *t_r_ ∈ T_r_* and noise-free structural density maps sampled from the clean domain *v_r_ ∈ V_r_*, the DUAL framework learns to denoise the experimental tomograms *t_r_*to the structural density map level *v_f_ ∈ V_f_* and to simulate synthetic tomograms *t_f_ ∈ T_f_* from *V_r_*. We assume there are two types of noises in the simulation branch: random non-structural noises and structure-related noises such as missing wedge effect, defocus, and spherical aberration. The cyclic structure of DUAL consists of a denoising branch with denoiser *Dn* and a simulation branch with generators *G_s_* to generate structural noise and *G_n_*to generate non-structural noise. The loss functions ensure that the generated synthetic tomograms (noisy domain) *T_f_* and structural density maps (clean domain) *V_f_* contain the essential structural information and be indiscriminable from real ones. Now, we introduce the denoising branch, the simulation branch, and the loss functions of DUAL in greater detail.

### Denoising branch

The denoising branch includes a denoiser *Dn* to translate an experimental tomogram to a noise-free structural density map. When applied to an experimental tomogram *t_r_*, the denoiser outputs fake structural density map: *v_f_* = *Dn*(*t_r_*). When applied to fake tomograms *t_f_* generated in the training process, the denoiser outputs 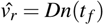, to reconstruct the input real density map *v_r_*.

### Simulation branch

The simulation branch includes two generators *G_s_* and *G_n_* and a noise encoder *E* to translate a noise-free structural density map to a noisy synthetic tomogram. When applied to a real structural density map *v_r_*, we first generate non-structural noise to distort *v_r_*. The non-structural noise is defined as purely random noises not related to the underlying structure. The non-structural noise *G_n_*(*z*) is generated by generator *G_n_*, where *z* is a random noise code sampled from a multivariate Gaussian distribution *N*(0, *I_K_*). We note that because of the highly non-linear nature of neural network *G_n_*, the output non-structural noise *G_n_* is not necessarily Gaussian. Assuming that the non-structural noises are independent for different voxels, we randomly permute (denoted by *P*) the non-structural noise mask generated by *G_n_*. Then, the synthetic tomogram is generated using the structural noise decoder *G_s_*:

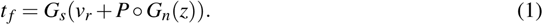

When applied to fake structural density maps *v_f_* generated in the training process, we first need to learn the noise pattern in the input experimental tomograms *t_r_* in order to reconstruct it. This is done by the encoder *E*. With the learned noise code *E*(*t_r_*), we apply *G_n_* and *G_s_* in the same way to reconstruct *t_r_*:

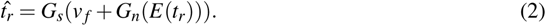

### Loss functions

DUAL is trained with three loss functions with different purposes. The reconstruction loss function ensures that the essential structural information is preserved in both branches and the noise pattern is properly learned from experimental tomogram *t_r_*. The adversarial loss function ensures that the generated *v_f_* and *t_f_* are indiscriminable from real ones *t_r_*and *v_r_* in style, respectively. The noise code regularization loss function ensures that the extracted noise code from *t_r_* follows a standard multivariate Gaussian distribution. The overall loss function is a linear combination of the three types of loss functions with weight coefficients λs:

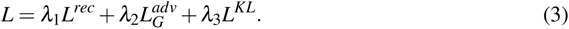

#### Reconstruction loss

The main idea behind the reconstruction loss function is that if the essential structural features are well-preserved by the simulation branch, the denoising branch can successfully bring back *v_r_* from *t_f_*. Similarly, if the essential structural features are well-preserved by the denoising branch and the noise pattern is properly encoded by the noise code *z*, the simulation branch can successfully bring back *t_r_* from *v_f_* and *z*.

To enforce the denoiser *Dn* to learn how to remove non-structural and structural noise from experimental tomograms, we minimize the difference between *v_r_* and 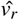, so as to maximize the consistency between real and reconstructed structural density maps. Specifically, we choose the mean squared error and Pearson’s correlation coefficient as the measure of the difference between *v_r_* and 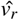. The reconstruction loss function for density maps is defined as:

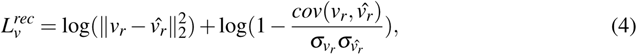

where *cov*() is the covariance function and σ denotes the standard deviation, to minimize the Euclidean distance and maximize the correlation.

Similarly, for the simulation branch, we minimize the ℓ_2_ loss and maximize the correlation between experimental tomogram *v_r_* and the reconstructed one 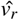. It is expected that the encoder *E* can learn to extract and encode non-structural noise information effectively. Therefore, when the noise code extracted from *v_r_* is decoded and added back in the simulation branch, the 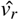 is expected to reconstruct *v_r_* with the correct noise pattern. The reconstruction loss function for tomograms is defined as:

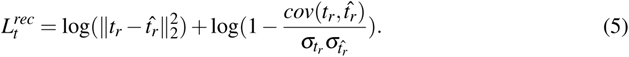

We combine them with equal weights to get the overall reconstruction loss function: 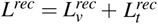.

#### Adversarial loss

In order to generate fake images *t_f_* and *v_f_* indistinguishable from real ones *t_r_*and *v_r_*, we train discriminators to discriminate them and then train generators *G_s_* and *G_n_*, encoder *E*, and denoiser *Dn* to minimize the adversarial loss function [57] from the discriminators. Because we do not have the corresponding ground truth for *v_f_* and *t_f_*, we introduce discriminators and adversarial loss functions to evaluate their similarity to *v_r_*and *t_f_* in style. To guide the discriminator for the noisy domain *D_t_* to assign higher scores to *t_r_*and lower scores to *t_f_*, we define the adversarial loss function for training *D_t_* as:

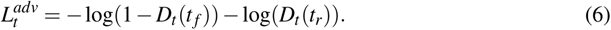

Similarly, the adversarial loss function for training the discriminator for the noisy domain *D_v_*is defined as:

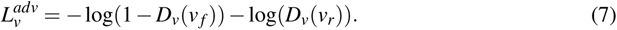

The combined loss function for training the discriminators is: 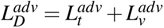.

After training the discriminators to classify real and fake images in each domain, we utilize them to improve the quality of *t_f_* and *v_f_* from generators and denoiser. To generate indistinguishable fake images, it is expected to increase the scores of *t_f_* and *v_f_* assigned by the discriminators. As a result, the adversarial loss function for training generators *G_s_* and *G_n_*, encoder *E*, and denoiser *Dn* is defined as:

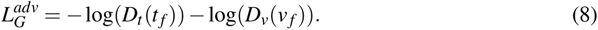

In each training iteration, the 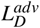 and 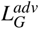 are optimized in an alternative manner.

#### Noise code regularization loss

There are two sources of noise codes: when reconstructing 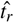, the noise code comes from encoder *E*(*v_r_*); while generating synthetic tomogram *t_f_*, we sample the noise code from a standard multi-variate Gaussian distribution *N*(0, *I_K_*). As the non-structural noise is generated from the noise code, generator *G_n_*may produce non-structural noises with different patterns for these two heterogeneous sources of noise codes. To overcome this issue, we introduce a noise code regularization loss function that aims to align the distributions of these two sources of noise codes.

To unify them, we enforce the noise codes from the encoder to follow a standard multivariate Gaussian distribution *N*(0, *I_K_*). The Kullback–Leibler divergence loss on the two distributions is defined as:

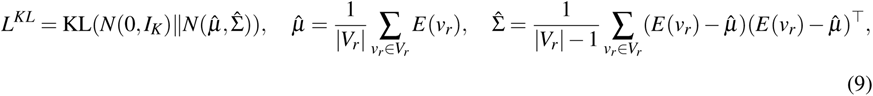

where 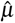 and 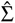 are estimated mean and covariance matrix from *E*(*V_r_*), extracted noise code from a training batch of samples from the noisy domain.

### Denoising quantitative measures

To evaluate the denoising performance, we choose two criteria, namely Peak Signal-to-Noise Ratio (PSNR) and Structural Similarity Index Measure (SSIM). For a reconstructed image 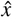 and its ground truth *x*, PSNR is defined based on mean squared error [63]:

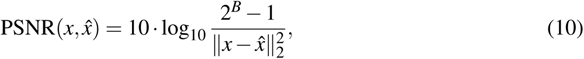

where *B* represents the number of bits for each pixel to be stored.

SSIM [64] is another important criterion for measuring imaging restoration quality. It is defined as:

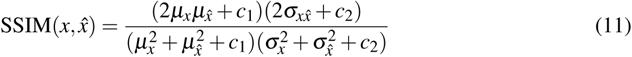

where 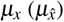 and 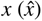 are the mean and variance of 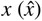. 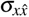 is covariance between 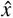 and 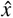. *c*_1_ and *c*_2_ are two constants to avoid instability brought by extremely small denominator values.

### Datasets and training preparation

SHREC 2021 benchmark: SHREC 2021 track: classification in cryo-electron tomograms [23] provides a synthetic cryo-ET benchmark dataset that consists of ten tomograms. Each tomogram corresponds to a noise-free grand model of structural density map of the same size. Each grand model contains randomly distributed vesicles (membrane structure), fiducial markers, and thirteen types of macromolecular complexes: TRiC (PDB ID: 4V94), 26S proteasome (4CR2), ClpB (1QVR), rubisco (1BXN), P97/vcp (3CF3), Cand1-Cul1-Roc1 (1U6G), Sse1p, Hsp70 (3D2F), Hsp90-Sba1 (2CG9), GET3 (3H84), Ssb1, Hsp70 (3GL1), LJ0536 S106A (3QM1), Hsp70 ATPase (1S3X), and yeast mito ribosome (5MRC). We split this dataset into three parts in an unpaired setting. The first four tomograms are used to construct the noisy domain of tomograms. The grand model of the next four tomograms are used to construct the clean domain of structural density maps. Similar to the SHREC 2021 track [23], the last tomogram and grand model pair is used as the testing dataset for evaluation.

Ribosome benchmark: this is a single-particle benchmark dataset to evaluate the subtomogram averaging performance of *RELION* [24]. A total of seven tomograms in this dataset contain purified 80S ribosomes from *Saccharomyces cerevisiae*. Individual ribosome locations are provided by the authors through manual picking. The original tomograms in this dataset are of voxel spacing 0.227 nm and tilt-angle range ±60°.

*Rattus* neuron dataset: this dataset contains six cellular tomograms from primary *Rattus* neuron culture [59]. Three types of macromolecular complexes: 26S proteasome, TRiC/CCT chaperonin, and ribosome, are detected and recovered by the authors through manual picking and subtomogram averaging. The original tomograms in this dataset are of voxel spacing 1.368 nm and tilt-angle range −50° to +70°.

*Chlamydomonas* pyrenoid dataset: this dataset contains one tomogram of the *Chlamydomonas reinhardtii* pyrenoid with abundant rubisco holoenzymes [11]. The original tomogram in this dataset is of voxel spacing 1.368 nm and tilt-angle range ±60°.

*Chlamydomonas* chloroplast dataset: this dataset contains four tomogram of the *Chlamydomonas reinhardtii* chloroplast [65]. Compared with other experimental datasets, the tomograms are of higher SNR due to their use of advanced direct detector cameras and the contrast-enhancing Volta phase plate. The original tomograms in this dataset are of voxel spacing 1.368 nm and tilt-angle range ±60°.

SARS-CoV-2 infection dataset: this dataset contains three tomograms of human airway epithelium infected by SARS-CoV-2 B.1.1.7 variant [66]. This dataset is collected under a conventional transmission electron microscope with a relatively high SNR. The original tomograms in this dataset are of voxel spacing 0.457 nm and dual-axis tilt-angle range ±60°.

Because the experimental datasets do not have available corresponding structural density maps, the four grand models the SHREC 2021 benchmark dataset is also used as the clean domain for experimental datasets. As the clean domain from the SHREC 2021 benchmark dataset has voxel spacing of 1 nm, we rescaled the voxel spacing of all tomograms in the five experimental datasets to 1 nm. Then, we standardized each tomogram or grand model by subtracting its mean and dividing by its standard deviation. To reduce memory consumption and increase the efficiency of neural network training, we divide the tomograms and grand models into non-overlapping subvolumes of size 32^2^ as inputs. We note that subvolumes of other sizes can also be processed. The larger the subvolume size, the fewer inputs need to be processed but also slower training speed for each sample. Training batches of samples were randomly selected and matched from the clean domain and noisy domain. When predicting on testing datasets using the trained neural networks of DUAL, we employed the overlap-tile strategy [56] to avoid artifacts at the boundary between subvolumes.

### Template-free particle picking

We applied a popular template-free particle picking algorithm Difference of Gaussians (DoG) [67]. DoG picks potential particles by detecting local maxima in the substraction of two Gaussian filtered versions of the tomogram with different standard deviations. We chose σ_1_ as 8.0 and σ_2_ as 8.8 with a multiplication factor *k* of 1.1. Overlapping detected local maxima within 24 nm of pairwise distances were filtered. Then, the top 500 DoG detections were selected for each tomogram and their denoised versions by each method.

### Implementation details

DUAL is implemented using *PyTorch* [68] with four Nvidia RTX 2080Ti GPU instances support. We chose AdamW [69] as the optimizer for all networks with a learning rate of 10*^−^*^4^, β_1_ of 0.9, β_2_ of 0.999, ε of 10*^−^*^8^ and weight decay of 10*^−^*^6^. For each dataset, the model was randomly initialized with an orthogonal kernel weight initializer and trained for 20 epochs. Loss coefficients were set empirically based on the performance as λ_1_ = 1, λ_2_ = 1, and λ_3_ = 10*^−^*^1^. The training algorithm is shown in supplementary note 4.

## Data source

The SHREC 2021 benchmark dataset is obtained from [23]. The ribosome benchmark dataset is obtained from EMPIAR 10045 [24]. The *Rattus* neuron dataset is obtained from [59]. The *Chlamydomonas* pyrenoid dataset is obtained from EMD-12749 [11]. The *Chlamydomonas* chloroplast dataset is obtained from EMD-10780 to EMD-10783 [65]. The SARS-CoV-2 infection dataset is obtained from EMD-14364, EMD-14365, and EMD-14367 [66].

## Code availability

To directly benefit the cryo-ET research community, we will disseminate all the code into our open-source cryo-ET data analysis software *AITom* [25]. Currently, we have disseminated 25 of our existing published algorithms into AITom. There are more than 20 tutorials provided in AITom for different cryo-ET analysis tasks with more than 30,000 lines of codes mainly written in python and C++. We will also integrate our code with the software *Scipion* [70] as a plugin. User-friendly tutorials will be provided on how to apply our models to users’ own datasets.

## Data availability

We will disseminate the subtomogram averages into EM Data Bank [71]. The trained models and demo data will be disseminated into *AITom* [25].

## Acknowledgements

This work was supported in part by U.S. NIH grants R01GM134020 and P41GM103712, and NSF grants DBI-1949629, DBI-2238093, IIS-2007595, IIS-2211597, and MCB-2205148. The computational resources were supported by AMD HPC Fund and by Dr. Zachary Freyberg’s lab. This work was supported in part by Oracle Cloud credits and related resources provided by Oracle for Research. X.Z and M.R.U. were supported in part by a fellowship from CMLH. We thank Dr. Qiang Guo for providing testing datasets, Michael Maxwell and Dr. Xingjian Li for critical comments on the manuscript.

## Author contributions

M.X. and X.Z. conceived the research. X.Z., and Y.D. designed the method. Y.D. and Y.Z. implemented the method. M.U. and A.D. gave suggestions. X.Z. refined the method and conducted the experiments. Y.D. and Y.Z. conducted the baseline experiments. X.Z., Y.D., and M.X. wrote the manuscript. All authors edited the manuscript.

## References

1. Miroslava Schaffer, Stefan Pfeffer, Julia Mahamid, Stephan Kleindiek, Tim Laugks, Sahradha Albert, Benjamin D Engel, Andreas Rummel, Andrew J Smith, Wolfgang Baumeister, et al. A cryo-fib lift-out technique enables molecular-resolution cryo-et within native caenorhabditis elegans tissue. Nature methods, 16(8):757–762, 2019.

2. Meijing Li, Jianfei Ma, Xueming Li, and Sen-Fang Sui. In situ cryo-et structure of phycobilisome–photosystem ii supercomplex from red alga. Elife, 10:e69635, 2021.

3. Amanda E Ward, Kelly A Dryden, Lukas K Tamm, and Barbie K Ganser-Pornillos. Catching hiv in the act of fusion: Insight from cryo-et intermediates of hiv membrane fusion. Biophysical Journal, 116(3):180a, 2019.

4. Antonio Martinez-Sanchez, Wolfgang Baumeister, and Vladan Lučić. Statistical spatial analysis for cryo-electron tomography. Computer Methods and Programs in Biomedicine, page 106693, 2022.

5. Yuewei Wang, Tong Huo, Yu-Jung Tseng, Lan Dang, Zhili Yu, Wenjuan Yu, Zachary Foulks, Rebecca L Murdaugh, Steven J Ludtke, Daisuke Nakada, et al. Using cryo-et to distinguish platelets during pre-acute myeloid leukemia from steady state hematopoiesis. Communications biology, 5(1):1–9, 2022.

6. Rui Wang, Rebecca L Stone, Jason T Kaelber, Ryan H Rochat, Alpa M Nick, K Vinod Vijayan, Vahid Afshar-Kharghan, Michael F Schmid, Jing-Fei Dong, Anil K Sood, et al. Electron cryotomography reveals ultrastructure alterations in platelets from patients with ovarian cancer. Proceedings of the National Academy of Sciences, 112(46):14266–14271, 2015.

7. Stephanie E Siegmund, Robert Grassucci, Stephen D Carter, Emanuele Barca, Zachary J Farino, Martí Juanola-Falgarona, Peijun Zhang, Kurenai Tanji, Michio Hirano, Eric A Schon, et al. Three-dimensional analysis of mitochondrial crista ultrastructure in a patient with leigh syndrome by in situ cryoelectron tomography. iScience, 6:83–91, 2018.

8. Yanhe Zhao, Justine Pinskey, Jianfeng Lin, Weining Yin, Patrick R Sears, Leigh A Daniels, Maimoona A Zariwala, Michael R Knowles, Lawrence E Ostrowski, and Daniela Nicastro. Structural insights into the cause of human rsph4a primary ciliary dyskinesia. Molecular biology of the cell, 32(12):1202–1209, 2021.

9. Xueming Li. Cryo-electron tomography: observing the cell at the atomic level. Nature Methods, 18(5):440–441, 2021.

10. Shoh Asano, Benjamin D Engel, and Wolfgang Baumeister. In situ cryo-electron tomography: a post-reductionist approach to structural biology. Journal of molecular biology, 428(2):332– 343, 2016.

11. Emmanuel Moebel, Antonio Martinez-Sanchez, Lorenz Lamm, Ricardo D Righetto, Wojciech Wietrzynski, Sahradha Albert, Damien Larivière, Eric Fourmentin, Stefan Pfeffer, Julio Ortiz, et al. Deep learning improves macromolecule identification in 3d cellular cryo-electron tomograms. Nature Methods, 18(11):1386–1394, 2021.

12. Muyuan Chen, Wei Dai, Stella Y Sun, Darius Jonasch, Cynthia Y He, Michael F Schmid, Wah Chiu, and Steven J Ludtke. Convolutional neural networks for automated annotation of cellular cryo-electron tomograms. nature methods, 14(10):983, 2017.

13. Chengqian Che, Ruogu Lin, Xiangrui Zeng, Karim Elmaaroufi, John Galeotti, and Min Xu. Improved deep learning-based macromolecules structure classification from electron cryotomograms. Machine Vision and Applications, 29(8):1227–1236, 2018.

14. Ran Li, Xiangrui Zeng, Stephanie E Sigmund, Ruogu Lin, Bo Zhou, Chang Liu, Kaiwen Wang, Rui Jiang, Zachary Freyberg, Hairong Lv, et al. Automatic localization and identification of mitochondria in cellular electron cryo-tomography using faster-rcnn. BMC bioinformatics, 20(3):75–85, 2019.

15. Ngan Nguyen, Ciril Bohak, Dominik Engel, Peter Mindek, Ondřej Strnad, Peter Wonka, Sai Li, Timo Ropinski, and Ivan Viola. Finding nano-\” otzi: Semi-supervised volume visualization for cryo-electron tomography. *arXiv preprint arXiv:2104.01554*, 2021.

16. Tristan Bepler, Kotaro Kelley, Alex J Noble, and Bonnie Berger. Topaz-denoise: general deep denoising models for cryoem and cryoet. Nature communications, 11(1):1–12, 2020.

17. Zhidong Yang, Fa Zhang, and Renmin Han. Self-supervised cryo-electron tomography volumetric image restoration from single noisy volume with sparsity constraint. In Proceedings of the IEEE/CVF International Conference on Computer Vision, pages 4056–4065, 2021.

18. Xiangrui Zeng and Min Xu. Gum-net: Unsupervised geometric matching for fast and accurate 3d subtomogram image alignment and averaging. In Proceedings of the IEEE/CVF Conference on Computer Vision and Pattern Recognition, pages 4073–4084, 2020.

19. Xiangrui Zeng, Gregory Howe, and Min Xu. End-to-end robust joint unsupervised image alignment and clustering. In Proceedings of the IEEE/CVF International Conference on Computer Vision, pages 3854–3866, 2021.

20. Xiangrui Zeng, Anson Kahng, Liang Xue, Julia Mahamid, Yi-Wei Chang, and Min Xu. Disca: high-throughput cryo-et structural pattern mining by deep unsupervised clustering. bioRxiv, 2021.

21. Pietro Perona and Jitendra Malik. Scale-space and edge detection using anisotropic diffusion. IEEE Transactions on pattern analysis and machine intelligence, 12(7):629–639, 1990.

22. Antoni Buades, Bartomeu Coll, and J-M Morel. A non-local algorithm for image denoising. In 2005 IEEE computer society conference on computer vision and pattern recognition (CVPR’05), volume 2, pages 60–65. Ieee, 2005.

23. Ilja Gubins, Marten L. Chaillet, Gijs van der Schot, M. Cristina Trueba, Remco C. Veltkamp, Friedrich Förster, Xiao Wang, Daisuke Kihara, Emmanuel Moebel, Nguyen P. Nguyen, Tommi White, Filiz Bunyak, Giorgos Papoulias, Stavros Gerolymatos, Evangelia I. Zacharaki, Konstantinos Moustakas, Xiangrui Zeng, Sinuo Liu, Min Xu, Yaoyu Wang, Cheng Chen, Xuefeng Cui, and Fa Zhang. SHREC 2021: Classification in Cryo-electron Tomograms. In Silvia Biasotti, Roberto M. Dyke, Yukun Lai, Paul L. Rosin, and Remco C. Veltkamp, editors, Eurographics Workshop on 3D Object Retrieval. The Eurographics Association, 2021.

24. Tanmay AM Bharat and Sjors HW Scheres. Resolving macromolecular structures from electron cryo-tomography data using subtomogram averaging in relion. Nature protocols, 11(11):2054– 2065, 2016.

25. Xiangrui Zeng and Min Xu. Aitom: Open-source ai platform for cryo-electron tomography data analysis. *arXiv preprint arXiv*:1911.03044, 2019.

26. Martin Turk and Wolfgang Baumeister. The promise and the challenges of cryo-electron tomography. FEBS letters, 594(20):3243–3261, 2020.

27. David N Mastronarde and Susannah R Held. Automated tilt series alignment and tomographic reconstruction in imod. Journal of structural biology, 197(2):102–113, 2017.

28. Jose-Jesus Fernandez and Sam Li. Tomoalign: A novel approach to correcting sample motion and 3d ctf in cryoet. Journal of Structural Biology, 213(4):107778, 2021.

29. Rui Yan, Singanallur V Venkatakrishnan, Jun Liu, Charles A Bouman, and Wen Jiang. Mbir: A cryo-et 3d reconstruction method that effectively minimizes missing wedge artifacts and restores missing information. Journal of structural biology, 206(2):183–192, 2019.

30. Yuchen Deng, Yu Chen, Yan Zhang, Shengliu Wang, Fa Zhang, and Fei Sun. Icon: 3d reconstruction with ‘missing-information’restoration in biological electron tomography. Journal of structural biology, 195(1):100–112, 2016.

31. Xinrui Huang, Sha Li, and Song Gao. Exploring an optimal wavelet-based filter for cryo-et imaging. Scientific reports, 8(1):1–9, 2018.

32. Emmanuel Moebel and Charles Kervrann. A monte carlo framework for denoising and missing wedge reconstruction in cryo-electron tomography. In International Workshop on Patch-based Techniques in Medical Imaging, pages 28–35. Springer, 2018.

33. Jochen Böhm, Achilleas S Frangakis, Reiner Hegerl, Stephan Nickell, Dieter Typke, and Wolfgang Baumeister. Toward detecting and identifying macromolecules in a cellular context: template matching applied to electron tomograms. Proceedings of the National Academy of Sciences, 97(26):14245–14250, 2000.

34. Antonio Martinez-Sanchez, Inmaculada Garcia, Shoh Asano, Vladan Lucic, and Jose-Jesus Fernandez. Robust membrane detection based on tensor voting for electron tomography. Journal of structural biology, 186(1):49–61, 2014.

35. Fernando Amat, Luis R Comolli, Farshid Moussavi, John Smit, Kenneth H Downing, and Mark Horowitz. Subtomogram alignment by adaptive fourier coefficient thresholding. Journal of structural biology, 171(3):332–344, 2010.

36. Min Xu, Martin Beck, and Frank Alber. High-throughput subtomogram alignment and classification by fourier space constrained fast volumetric matching. Journal of structural biology, 178(2):152–164, 2012.

37. Benjamin A Himes and Peijun Zhang. emclarity: software for high-resolution cryo-electron tomography and subtomogram averaging. Nature methods, 15(11):955, 2018.

38. Paula P Navarro, Henning Stahlberg, and Daniel Castaño-Díez. Protocols for subtomogram averaging of membrane proteins in the dynamo software package. Frontiers in molecular biosciences, page 82, 2018.

39. Sjors HW Scheres. Relion: implementation of a bayesian approach to cryo-em structure determination. Journal of structural biology, 180(3):519–530, 2012.

40. Mohamad Harastani, Mikhail Eltsov, Amélie Leforestier, and Slavica Jonic. Hemnma-3d: Cryo electron tomography method based on normal mode analysis to study continuous conformational variability of macromolecular complexes. Frontiers in molecular biosciences, 8, 2021.

41. Mohamad Harastani, Mikhail Eltsov, Amélie Leforestier, and Slavica Jonic. Tomoflow: Analysis of continuous conformational variability of macromolecules in cryogenic subtomograms based on 3d dense optical flow. Journal of molecular biology, 434(2):167381, 2022.

42. Sinuo Liu, Yan Ma, Xiaojuan Ban, Xiangrui Zeng, Vamsi Nallapareddy, Ajinkya Chaudhari, and Min Xu. Efficient cryo-electron tomogram simulation of macromolecular crowding with application to sars-cov-2. In 2020 IEEE International Conference on Bioinformatics and Biomedicine (BIBM), pages 80–87. IEEE, 2020.

43. Guang Tang, Liwei Peng, Philip R Baldwin, Deepinder S Mann, Wen Jiang, Ian Rees, and Steven J Ludtke. Eman2: an extensible image processing suite for electron microscopy. Journal of structural biology, 157(1):38–46, 2007.

44. Niall O’Mahony, Sean Campbell, Anderson Carvalho, Suman Harapanahalli, Gustavo Velasco Hernandez, Lenka Krpalkova, Daniel Riordan, and Joseph Walsh. Deep learning vs. traditional computer vision. In Science and information conference, pages 128–144. Springer, 2019.

45. Long Pei, Min Xu, Zachary Frazier, and Frank Alber. Simulating cryo electron tomograms of crowded cell cytoplasm for assessment of automated particle picking. BMC bioinformatics, 17(1):405, 2016.

46. Ilja Gubins, Gijs van der Schot, Remco C Veltkamp, FG Förster, Xuefeng Du, Xiangrui Zeng, Zhenxi Zhu, Lufan Chang, Min Xu, Emmanuel Moebel, et al. Classification in cryo-electron tomograms. SHREC’19 Track, 2019.

47. Peter Scheible, Salim Sazzed, Jing He, and Willy Wriggers. Tomosim: Simulation of filamentous cryo-electron tomograms. In 2021 IEEE International Conference on Bioinformatics and Biomedicine (BIBM), pages 2560–2565. IEEE, 2021.

48. Lorenz Lamm, Ricardo D Righetto, Wojciech Wietrzynski, Matthias Pöge, Antonio Martinez-Sanchez, Tingying Peng, and Benjamin D Engel. Membrain: A deep learning-aided pipeline for automated detection of membrane proteins in cryo-electron tomograms. bioRxiv, 2022.

49. Charith A Hettiarachchi, Matthew T Swulius, and Federico M Harte. Assessing constituent volumes and morphology of bovine casein micelles using cryo-electron tomography. Journal of dairy science, 103(5):3971–3979, 2020.

50. Mirabela Rusu, Zbigniew Starosolski, Manuel Wahle, Alexander Rigort, and Willy Wriggers. Automated tracing of filaments in 3d electron tomography reconstructions using sculptor and situs. Journal of structural biology, 178(2):121–128, 2012.

51. Emmanuel Moebel and Charles Kervrann. A monte carlo framework for missing wedge restoration and noise removal in cryo-electron tomography. Journal of Structural Biology: X, 4:100013, 2020.

52. Haonan Zhang, Yan Li, Yanan Liu, Dongyu Li, Lin Wang, Kai Song, Keyan Bao, and Ping Zhu. Rest: A method for restoring signals and revealing individual macromolecule states in cryo-et. bioRxiv, 2022.

53. Jun-Yan Zhu, Taesung Park, Phillip Isola, and Alexei A Efros. Unpaired image-to-image translation using cycle-consistent adversarial networks. In Proceedings of the IEEE international conference on computer vision, pages 2223–2232, 2017.

54. Sinuo Liu, Xiaojuan Ban, Xiangrui Zeng, Fengnian Zhao, Yuan Gao, Wenjie Wu, Hongpan Zhang, Feiyang Chen, Thomas Hall, Xin Gao, et al. A unified framework for packing deformable and non-deformable subcellular structures in crowded cryo-electron tomogram simulation. BMC bioinformatics, 21(1):1–24, 2020.

55. Aziz Alotaibi. Deep generative adversarial networks for image-to-image translation: A review. Symmetry, 12(10):1705, 2020.

56. Olaf Ronneberger, Philipp Fischer, and Thomas Brox. U-net: Convolutional networks for biomedical image segmentation. In International Conference on Medical image computing and computer-assisted intervention, pages 234–241. Springer, 2015.

57. Ian Goodfellow, Jean Pouget-Abadie, Mehdi Mirza, Bing Xu, David Warde-Farley, Sherjil Ozair, Aaron Courville, and Yoshua Bengio. Generative adversarial nets. Advances in neural information processing systems, 27, 2014.

58. Jaakko Lehtinen, Jacob Munkberg, Jon Hasselgren, Samuli Laine, Tero Karras, Miika Aittala, and Timo Aila. Noise2noise: Learning image restoration without clean data. *arXiv preprint arXiv:1803.04189*, 2018.

59. Qiang Guo, Carina Lehmer, Antonio Martínez-Sánchez, Till Rudack, Florian Beck, Hannelore Hartmann, Manuela Pérez-Berlanga, Frédéric Frottin, Mark S Hipp, F Ulrich Hartl, et al. In situ structure of neuronal c9orf72 poly-ga aggregates reveals proteasome recruitment. Cell, 172(4):696–705, 2018.

60. Cindi L Schwartz, Vasily I Sarbash, Fazoil I Ataullakhanov, J Richard Mcintosh, and Daniela Nicastro. Cryo-fluorescence microscopy facilitates correlations between light and cryo-electron microscopy and reduces the rate of photobleaching. Journal of microscopy, 227(2):98–109, 2007.

61. Xiaobo Zhai, Dongsheng Lei, Meng Zhang, Jianfang Liu, Hao Wu, Yadong Yu, Lei Zhang, and Gang Ren. Lottor: an algorithm for missing-wedge correction of the low-tilt tomographic 3d reconstruction of a single-molecule structure. Scientific reports, 10(1):1–17, 2020.

62. Yun-Tao Liu, Heng Zhang, Hui Wang, Chang-Lu Tao, Guo-Qiang Bi, and Z Hong Zhou. Isotropic reconstruction for electron tomography with deep learning. Nature communications, 13(1):1–17, 2022.

63. Alain Horé and Djemel Ziou. Is there a relationship between peak-signal-to-noise ratio and structural similarity index measure? IET Image Processing, 7(1):12–24, 2013.

64. Zhou Wang, Alan C Bovik, Hamid R Sheikh, and Eero P Simoncelli. Image quality assessment: from error visibility to structural similarity. IEEE transactions on image processing, 13(4):600– 612, 2004.

65. Wojciech Wietrzynski, Miroslava Schaffer, Dimitry Tegunov, Sahradha Albert, Atsuko Kanazawa, Jürgen M Plitzko, Wolfgang Baumeister, and Benjamin D Engel. Charting the native architecture of chlamydomonas thylakoid membranes with single-molecule precision. Elife, 9:e53740, 2020.

66. Andreia L Pinto, Ranjit K Rai, Jonathan C Brown, Paul Griffin, James R Edgar, Anand Shah, Aran Singanayagam, Claire Hogg, Wendy S Barclay, Clare E Futter, et al. Ultrastructural insight into sars-cov-2 entry and budding in human airway epithelium. Nature communications, 13(1):1–14, 2022.

67. NR Voss, CK Yoshioka, M Radermacher, CS Potter, and B Carragher. Dog picker and tiltpicker: software tools to facilitate particle selection in single particle electron microscopy. Journal of structural biology, 166(2):205–213, 2009.

68. Adam Paszke, Sam Gross, Francisco Massa, Adam Lerer, James Bradbury, Gregory Chanan, Trevor Killeen, Zeming Lin, Natalia Gimelshein, Luca Antiga, et al. Pytorch: An imperative style, high-performance deep learning library. Advances in neural information processing systems, 32, 2019.

69. Ilya Loshchilov and Frank Hutter. Decoupled weight decay regularization. *arXiv preprint arXiv:1711.05101*, 2017.

70. JM De la Rosa-Trevín, A Quintana, L Del Cano, A Zaldívar, I Foche, J Gutiérrez, J Gómez-Blanco, J Burguet-Castell, J Cuenca-Alba, V Abrishami, et al. Scipion: A software framework toward integration, reproducibility and validation in 3d electron microscopy. Journal of structural biology, 195(1):93–99, 2016.

71. Catherine L Lawson, Matthew L Baker, Christoph Best, Chunxiao Bi, Matthew Dougherty, Powei Feng, Glen Van Ginkel, Batsal Devkota, Ingvar Lagerstedt, Steven J Ludtke, et al. Emdatabank. org: unified data resource for cryoem. Nucleic acids research, 39(suppl_1):D456–D464, 2010.

